# HLH-3 Is Required for the Maturation of Neurons Necessary for Male Specific Exploration Behavior

**DOI:** 10.1101/2025.03.20.644401

**Authors:** Kimberly Goodwin, Aixa Alfonso

**Affiliations:** Biological Sciences, University of Illinois at Chicago, Chicago Illinois 60607, USA

**Keywords:** *hlh-3*, differentiation, AIM, ASJ, AWA, sexually dimorphic

## Abstract

Sexually dimorphic gene expression within the nervous system of many organisms gives rise to features that underline sexually dimorphic behaviors. In *C. elegans*, terminal differentiation of neurons with sexually dimorphic features often occur not at the time of neuronal birth, but rather during the juvenile to adult stage transition, when sexual maturation occurs. Previous works have shown that the bHLH transcription factor, HLH-3, is necessary for the differentiation of sex-specific neurons in hermaphrodites. Expression of the bHLH transcription factor, *hlh-3*, is dynamic and widespread throughout the embryo. Post-embryonic expression in the hermaphrodite is restricted to the HSNs, and the P cells and their descendants, including the VCs. In this work, we characterize the post-embryonic expression pattern of *hlh-3* in the male. Our findings show that the pattern of expression observed in males varies dramatically from that observed in hermaphrodites. Males show expanded expression in the head at the L1 and late L3/early L4 stage, as well as broader expression in the tail at the early L3/Late L4 stage. This leads us to the conclusion that the post-embryonic expression pattern of *hlh-3* is sexually dimorphic. We also describe the role of *hlh-3* in the maturation of a subset of neurons required for male specific exploration behavior. Our findings show that in the absence of *hlh-3* function, the AWA, ASJ, and AIM neurons fail to acquire terminally differentiated features and therefore contribute to the failure of males to perform behaviors mediated by these cells.

## INTRODUCTION

In multicellular organisms, neuronal tissue comprises a variety of specialized cell types which take part in regulatory circuitries that govern behavior. These different cell types acquire their morphology, neurotransmitter phenotype, and connectivity as a function of time and based on the transcriptomic profile that promotes their differentiation (Hobert, 2016; Tekieli et al., 2021). Sexually dimorphic neuronal maturation is controlled by two major processes, acquisition of sexual identity and developmental timing (H. Lawson et al., 2019). A complete cell lineage map, connectome, transcriptome, and a variety of transgenic reporters make the nematode *Caenorhabditis elegans* an ideal model organism to elucidate the mechanisms through which neurons obtain their identity (Cook et al., 2019; Sulston & Horvitz, 1977; The C. elegans Sequencing Consortium, 1998; Tintori et al., 2016).

Many sex shared neurons in *C. elegans* males have sexually dimorphic functions which control behaviors that support the drive to locate a mate and reproduce (Portman, 2017). One such behavior is sexually dimorphic exploration. This behavior is displayed by well fed, sexually mature, adult males which prioritize leaving a food source, increasing their likelihood of encountering a potential mate (Lipton et al., 2004; Ryan et al., 2014). This behavior is controlled by a network of neurons which sense internal signals that report reproductive drive and nutritional status (Barrios, 2014; Barrios et al., 2012; Hilbert & Kim, 2017, 2018; Ryan et al 2014; Lawson et. al., 2014; Sanzo-Machuca et al., 2019). The maturation of the sex shared neurons involved in this regulatory circuit takes place during the larval to adult transition (Hilbert & Kim, 2018; Ryan et al., 2014; Serrano-Saiz et al., 2017). Both developmental timing and sex identity have been implicated in the acquisition of terminal identity features like the downregulation of the diacetyl GPCR, ODR-10, in the AWAs, up-regulation of the TGF-ꞵ-like DAF-7 in the ASJs, and the neurotransmitter identity switch in the AIMs (Hilbert & Kim, 2018; H. Lawson et al., 2019; H. N. Lawson et al., 2020; Pereira et al., 2019; Ryan et al., 2014). Asymmetric expression of proneural genes in the descendants of ectodermal cells generate neuronal progenitor cells (Duggan & Chalfie, 1995; Ghysen & Dambly-Chaudiere, 1989). Most proneural genes encode proteins in the bHLH family (Bertrand et. al. 2002, Grove et. al. 2009). While proneural genes have been traditionally thought to have an early role promoting neuronal lineages, recent evidence suggests that some bHLH transcription factors have a direct influence on terminal fate acquisition too (Hobert, 2016; Masoudi et al., 2018; Poole et al., 2011). Previous findings implicate the bHLH (basic helix-loop-helix) transcription factor HLH-3 in the terminal differentiation of sex-specific neurons in *C. elegans* hermaphrodites (Doonan et al., 2008; Lloret-Fernández et al., 2018; Perez & Alfonso, 2020; Raut, 2017).

HLH-3 is a Class II bHLH transcription factor that heterodimerizes with the Class I bHLH transcription factor, HLH-2, to promote transcription of its targets (Grove et al., 2009; Krause et al., 1997). The gene *hlh-3* encodes a protein with 59% identity in the bHLH domain to the Asense proneural protein in Drosophila and 62% identity to MASH-1 in mice (Krause et. al. 1997, Doonan et. al. 2008). We have demonstrated that expression of *hlh-3* is required for the differentiation of ventral cord type C motor neurons (VCs) and hermaphrodite specific neurons (HSNs) (Doonan et al., 2008; Perez & Alfonso, 2020; Raut, 2017). Expression of the LIM homeobox transcription factor and terminal differentiation marker TTX-3 in the sex shared AIY neurons also requires HLH-3, however its function is redundant with two other bHLH proteins, HLH-16 and REF-2 (Filippopoulou et al., 2021; Murgan et al., 2015; F. Zhang et al., 2014). The role of HLH-3 extends beyond traditional proneural functions. For example, the sisters of a pair of neurosecretory cells, the NSMs, die because HLH-3 promotes expression of the apoptosis signal EGL-1 in them (Thellmann et al., 2003). Likewise, specification of the linker cell (LC), a migrating cell which leads the formation of the male somatic gonad, requires *hlh-3* function (Kimble & Hirsh, 1979; Sallee et al., 2017).

Extensive work has been done to characterize the expression of *hlh-3* in hermaphrodites (Doonan et al., 2008; Lloret-Fernández et al., 2018; Perez & Alfonso, 2020). Expression data was first gathered using a full-length translational fusion as well as a fosmid reporter, which showed similar patterns of expression (Doonan et al., 2008; Krause et al., 1997; Lloret-Fernández et al., 2018). This work showed that expression of *hlh-3* is broad and dynamic at the embryonic stage but becomes restricted to the HSNs and P cells upon hatching (Doonan et al., 2008; Krause et al., 1997; Lloret-Fernández et al., 2018). Further analysis showed widespread expression in the ring ganglia and ventral nerve cord (VNC). Expression of *hlh-3* in Pn.a (neural precursor) descendants in the L1 stage became restricted to VCs (Pn.aap) and HSNs by the L3 stage (Doonan et al., 2008). Generation and analysis of a CRISPR-Cas9 engineered allele, *hlh-3(ic271[ic271[hlh-3::gfp]])*, also displayed *hlh-3* expression restricted to HSNs and VCs after the L1 stage and throughout the lifetime of the hermaphrodite (Perez & Alfonso, 2020).

Previous efforts characterized the expression of *hlh-3* in hermaphrodite development. In this work we characterized the endogenous expression pattern of *hlh-3* in males throughout post-embryonic development utilizing the previously generated CRISPR-Cas9 allele, *hlh-3(ic271[hlh-3::gfp])*. We also investigated the role that HLH-3 plays in the maturation of sex shared neurons that regulate male specific exploration behavior.

## RESULTS

### Post-embryonic expression of *hlh-3* is sexually dimorphic

To characterize expression in the first larval stage (L1) of males, individual males were selected by identifying the large nuclei of Y and B cells in their tail (Sulston & Horvitz, 1977). In contrast to the pattern of expression observed in hermaphrodites, we find that *hlh-3* is expressed in several cells in the head of males at the L1 stage, these are not detectable in hermaphrodites (Figure 1 A and E). Like hermaphrodites, early L1 males show expression of *hlh-3* in 12 P cells. However, expression of *hlh-3* is seen in two additional cells in the mid body of hermaphrodites, the HSNs, which are absent in males (Figure 1 B and F). Expression in the HSNs was observed throughout the L1 stage of hermaphrodites. In the mid to late stage of both sexes, *hlh-3* expression remains present along the ventral cord (Figure 1 C, D, G and H).

**Figure 1:**
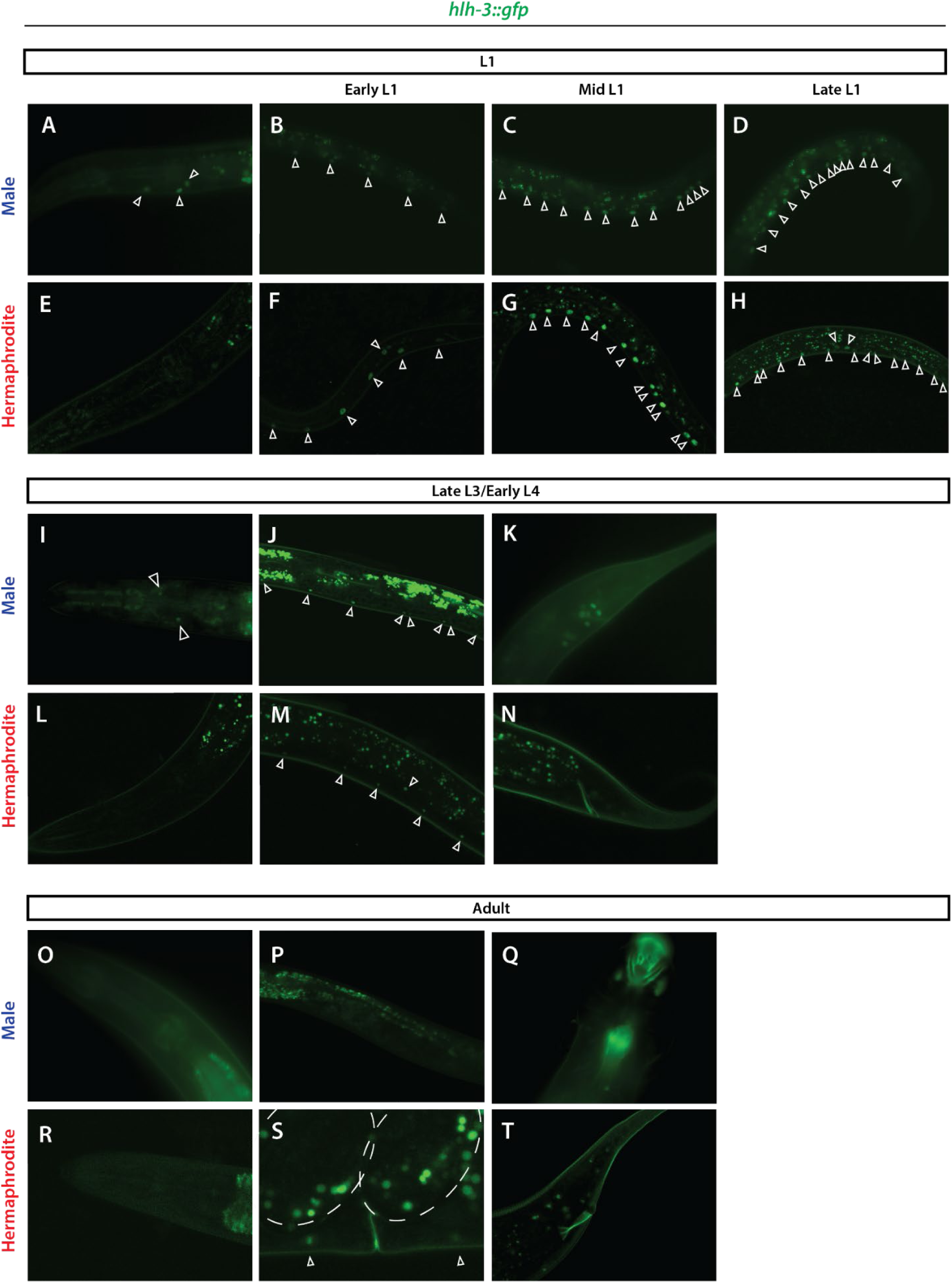
Post-embryonic expression of *ic271[hlh-3::gfp]* in males is sexually dimorphic. **(A**) Representative images of *ic271[hlh-3::gfp]* expression in the head of mid stage L1 male. **(B-D)** Representative images of midbody *ic271[hlh-3::gfp]* expression in early (∼16hrs post egg lay), mid (∼18hrs post egg lay), and late (∼20hrs post egg lay) stage L1 male ventral cord. **(E)** Representative images of *ic271[hlh-3::gfp]* expression in the head of mid stage L1 hermaphrodite **(F-H)** Representative images of midbody *ic271[hlh-3::gfp]* expression in early (∼16hrs post egg lay), mid (∼18hrs post egg lay), and late (∼20hrs post egg lay) stage L1 hermaphrodite ventral cord and midbody. Arrowheads point to positive nuclei, anterior is to the left, ventral is down. **(I-K)** Representative images of *ic271[hlh-3::gfp]* expression in head, midbody and tail of late L3/early L4 males, respectively. **(L-N)** Representative images of *ic271[hlh-3::gfp]* expression in head, midbody and ventral cord, and tail of late L3/early L4 hermaphrodites, respectively. Arrowheads point to positive nuclei, anterior is to the left, ventral is down. **(O-Q)** Representative images of *ic271[hlh-3::gfp]* expression in head, midbody and tail of adult males, respectively. Arrowheads point to positive nuclei. **(R-T)** Representative images of *ic271[hlh-3::gfp]* expression in head, ventral cord and tail of adult hermaphrodites, respectively. Dashed lines outline embryos. Arrowheads point to positive nuclei. Arrowheads point to positive nuclei, anterior is to the left, ventral is down.

Expression of *hlh-3* in males is broader than that in hermaphrodites. At the late L3/early L4 stages, positive cells in males included two cells in the head, several cells along the ventral cord, and a cluster at the tail ganglia (Figure 1 I-K). In contrast, positive cells in hermaphrodites are restricted to six cells along the ventral cord corresponding to the location of the VCs, and two cells in the midbody located between VC4 and VC5 corresponding to the location of the HSNs (Figure 1 L-N).

Expression of *hlh-3* in males is completely extinguished by the adult stage (Figure 1 O-Q). However, in adult hermaphrodites, expression was still observed in the VCs, as previously reported, but not in the head or tail (Figure 1 H, I, and J) (Perez & Alfonso, 2020).

### *hlh-3* null males do not exhibit exploration behavior

As published, loss of *hlh-3* function results in the hermaphrodite specific egg-laying defective (Egl) behavior (Doonan et al., 2008). Thus, we set out to determine if *hlh-3(tm1688)* adult males had any observable mating-related behavioral defects. A well characterized behavior specific to adult males is exploration (Lipton et al., 2004; Ryan et al., 2014). Well-fed, sexually mature adult males will leave a food source to explore, thus increasing their probability of encountering a potential mate (Lipton et al., 2004; Ryan et al., 2014). To address whether loss of *hlh-3* function affects this behavior, we performed a food leaving assay (Lipton et al., 2004). For these assays, we created a strain with the null *hlh-3(tm1688*) allele and a *him-5(e1490)* mutation to increase the production of males (Doonan et al., 2008; Hodgkin et al., 1979). We find that *hlh-3(tm1688);him-5(e1490)* adult males had a significantly lower probability to leave their food source when compared to strains containing only *him-5(e1490)* (Figure 2 A and B). This phenotype is not observed in *hlh-3(tm1688);him-5(e1490)* adult hermaphrodites. To address whether the lack of exploration resulted from a locomotion defect, we measured the ability of worms to reach the same distance as the leaving assay, when the entire area within the scorable distance is a lawn of bacteria (Barrios et al., 2012). We find that *hlh-3(tm1688);him-5(e1490)* mutant males can reach the edge of the food source as well as *him-5(e1490)* males do, thus a locomotion defect is not the cause of the observed lack of exploration (Figure S1 A and B). These results suggest that HLH-3 has a role in the differentiation of one or more neurons regulating the male mate searching drive (Barrios et al., 2012; Hilbert & Kim, 2017; Lipton et al., 2004; Ryan et al., 2014).

**Figure 2:**
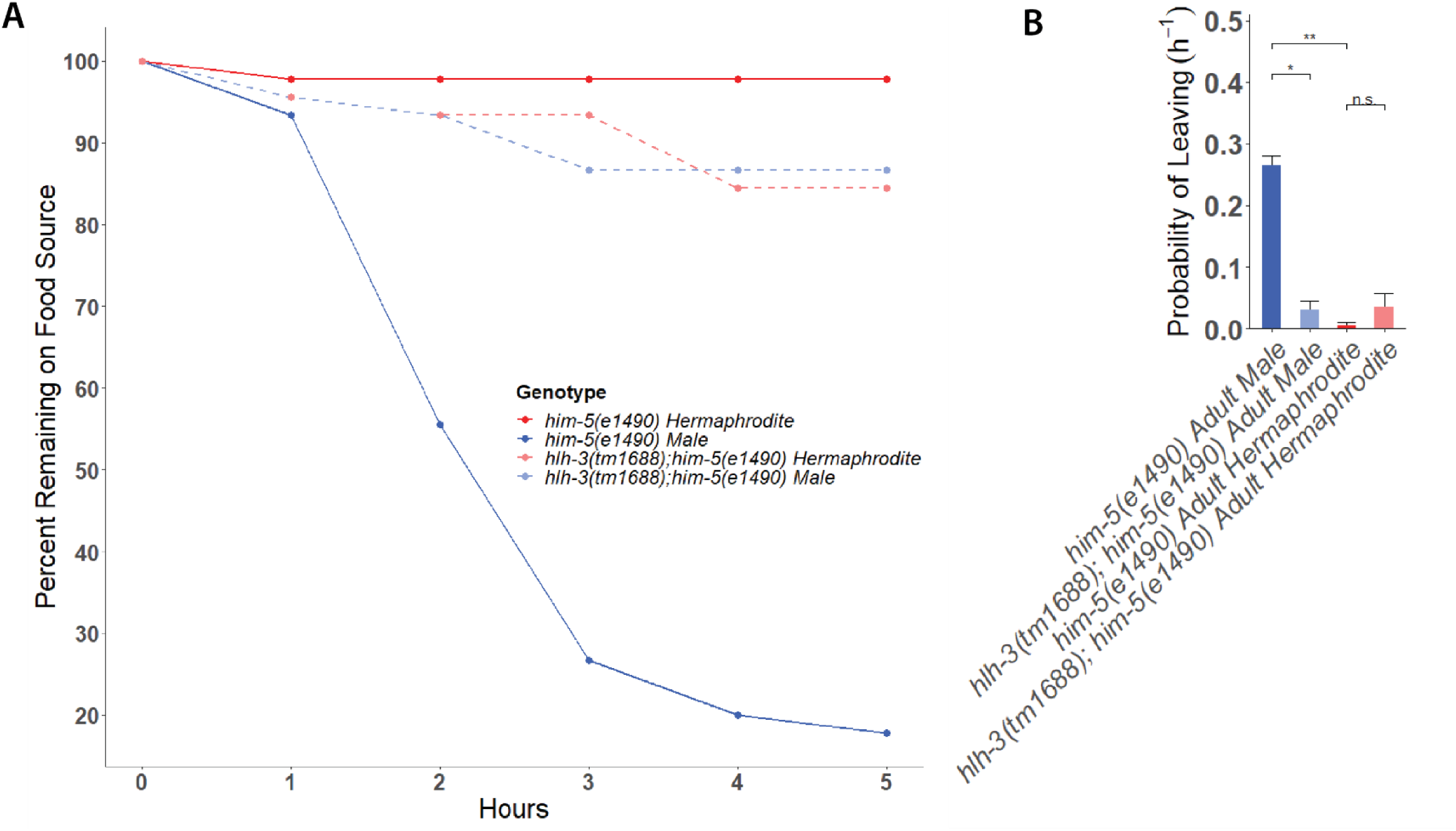
HLH-3 is required for male specific exploration behavior. **(A)** Representative graph depicting quantification of average male specific exploration behavior of *him-5(e1490)* adult males and *hlh-3(tm1688);him-5(e1490)* adult males (n = 45 for each genotype). Each point represents the percentage of worms remaining in the non-leaver area at the indicated time point. **(B)** Probability of leaving a food source for *him-5(e1490)* males and *hlh-3(tm1688);him-5(e1490)* adult males was calculated by the single exponential decay function N(t)/N(0)=exp(-λt), where N(t)/N(0) refers to the ratio of non-leavers to leavers at a given time (t), and the hazard value (λ) is the constant rate demonstrating the probability of leaving. Values plotted as averages with SEM for 3 independent experiments. *n.s. p = 0.961256* as determined by chi-squared analysis.

### Adult mutant male AIMs fail to fully switch their neurotransmitter identity

AIM interneurons are sex-shared neurons responsible for male mating drive and leaving behavior (Barrios et al., 2012). In males, adult AIM cells display a cholinergic identity. The neurotransmitter identity change from glutamatergic to cholinergic happens in the L4 to adult stage transition, a sexually dimorphic change (Pereira et al., 2015). Given that previous work has shown that *hlh-3(tm1688)* hermaphrodites show a lack of terminal differentiation in sex specific neurons, we wanted to determine if *hlh-3(tm1688)* adult male AIMs showed normal AIM sexual differentiation. To address whether adult mutant AIMs undergo maturation in the absence of *hlh-3* function, we asked whether *hlh-3(tm1688);him-5(e1490)* adult male AIMs attain their mature cholinergic neurotransmitter identity.

We assessed this by characterizing *eat-4*/VGLUT expression, a marker for glutamatergic identity, and *cho-1*/ChT expression, a marker for cholinergic identity (Lee et al., 1999; Matthies et al., 2006). We find that about 60% of adult *hlh-3(tm1688);him-5(e1490)* males retain expression of *eat-4*/VGLUT in AIM cells (Figure 3 A and C). To ascertain *cho-1*/ChT expression levels in AIMs, we measured intensity of expression relative to that of neighboring cholinergic AIY cells (Pereira et al., 2015). We find that AIMs in *hlh-3(tm1688); him-5(e1490)* adult males have a modest decrease in *cho-1*/ChT intensity when compared to the AIY cells in *him-5(e1490)* adult males, but there is no significant difference between that of *him-5(e1490)* and *hlh-3(tm1688);him-5(e1490)* L4 males (Figure 3 D-H). To determine whether this dimmer *cho-1*/ChT expression pattern in mutant adult males resembles that of *him-5(e1490)* L4 males we compared intensity of *hlh-3(tm1688);him-5(e1490)* adult male AIMs to that of *him-5(e1490)* L4 males. We find that while expression of *cho-1*/ChT decreases in *hlh-3(tm1688);him-5(e1490)* adult males, it is still significantly brighter than *him-5(e1490)* L4 males (Figure 3 H). Thus, while the AIMs in *hlh-3(tm1688);him-5(e1490)* adult males do not resemble those of *him-5(e1490)* L4 males, it is apparent they are not fully undergoing a neurotransmitter identity switch.

**Figure 3:**
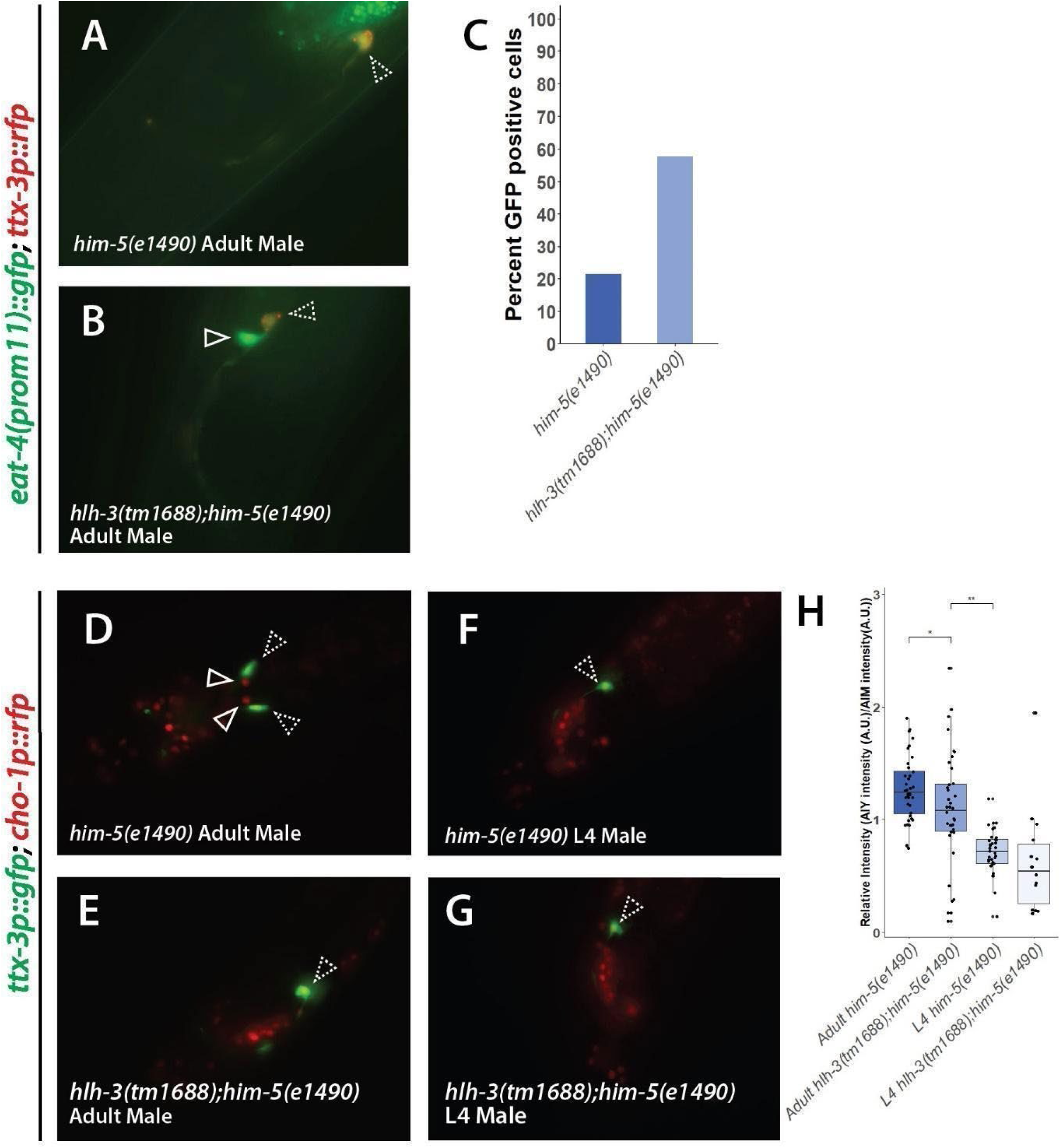
*hlh-3* is required for AIM terminal differentiation in adult males. **(A-B)** Representative images of *eat-4(prom11)::GFP* and *ttx-3p::RFP* expression in a *him -5(e1490)* and a *hlh-3(tm1688);him-5(e1490)* adult male, respectively. Open arrowheads indicate AIM cells, dashed arrowhead indicates AIY cells. **(C)** Percent of *eat-4(prom11)::GFP* positive AIM cells present in *him-5(e1490)* adult males (n = 42), and *hlh-3(tm1688);him-5(e1490)* adult males (n = 46). **(D-E)** Representative images of *cho-1p::RFP* and *ttx-3p::GFP* expression in a *him-5(e1490)* and a *hlh-3(tm1688);him-5(e1490)* adult male. Open arrowheads indicate AIM cells, dashed arrowhead indicates AIY cells. **(F-G)** Representative images of *cho-1p::RFP* and *ttx-3p::GFP* expression in a wild type and a *hlh-3(tm1688)* L4 male. Open arrowheads indicate AIM cells, dashed arrowhead indicates AIY cells. **(H)** Quantification of the relative intensity of *cho-1p::RFP* in AIM cells. Relative AIM intensity was determined by comparing its intensity to that of the standard *cho-1p::RFP* intensity in AIY cells of *him-5(e1490)* adult males (n = 36), *hlh-3(tm1688);him-5(e1490)* adult males (n = 38), *him-5(e1490)* L4 males (n = 33), and *hlh-3(tm1688);him-5(e1490)* L4 males (n =15). *\* p = 0.031623, ** p = 0.001101* as determined by Student’s T-test

### Mutant male ASJ sensory neurons do not express the TGF-β like *daf-7*

Exploration behavior in adult males is promoted by PDF-1 secretion via AIMs (Barrios et al., 2012). In males, reception of PDF-1 by the ASJ cells has been shown to increase *daf-7* expression and promote exploration (Hilbert & Kim, 2018). Due to the proximity of the two cells, it is possible that the secreted neuropeptide PDF-1 from AIM results in increased DAF-7/TGFß levels in ASJs of adult males (Cook et al., 2019). Since in *hlh-3(tm1688)* mutants, male AIMs are not terminally differentiated, then we expect to see a decrease in the levels of *daf-7* in males, but no measurable change in hermaphrodites. To address whether expression of *daf-7* was altered in *hlh-3(tm1688);him-5(e1490)* adult males, we examined *daf-7* expression using a *daf-7p::gfp* reporter (Hilbert ,& Kim, 2017). *daf-7* is expressed in one set of cells, the more anteriorly positioned ASIs, in juvenile males and hermaphrodites of all stages, and two sets of cells in adult males, the more anterior ASIs and the more posterior ASJs (Hilbert & Kim, 2017, Cook et al., 2019). Utilizing this information, we were able to identify the pattern of expression in *him-5(e1490)* and *hlh-3(tm1688);him-5(e1490)* strains. We find that expression of *daf-7* was absent in one or more ASJ cells in *hlh-3(tm1688); him-5(e1490)* adult males, whereas expression in another cell, the ASIs remain unaltered (Figure 4 A, B, E, and F) (Hilbert & Kim, 2017). Examination of *daf-7* in *hlh-3(tm1688);him-5(e1490)* adult hermaphrodites showed no change in expression pattern (Figure 4 C, D, E, and F).

**Figure 4:**
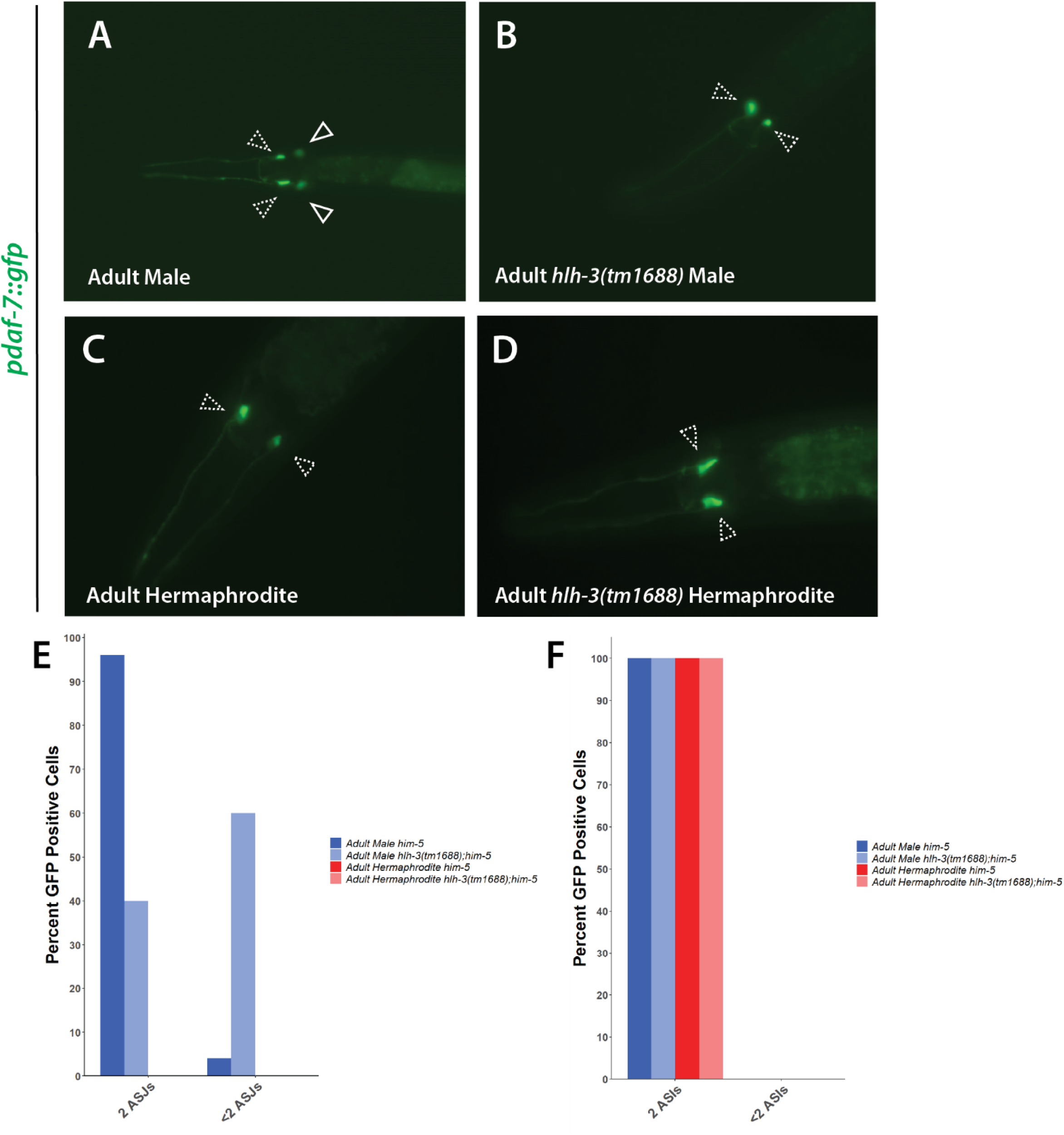
Upregulation of *daf-7* in adult male ASJs requires *hlh-3*. **(A-B)** Representative images of *daf-7p::gfp* expression in *him-5(e1490)* and *hlh-3(tm1688);him-5(e1490)* adult males. **(C-D)** Representative images of *daf-7p::gfp* expression in *him-5(e1490)* and *hlh-3(tm1688) him-5(e1490)* adult hermaphrodites. Dashed arrowheads indicate ASI cells, Solid arrowheads indicate ASJ cells. **(E)** Quantification of individuals of the indicated genotype and sex containing < 2 or 2 *daf-7p::gfp* positive ASJ and ASI cells for *him-5(e1490)* adult males (n = 26), *hlh-3(tm1688);him-5(e1490)* adult males (n = 30), *him-5(e1490)* hermaphrodites (n = 20), and *hlh-3(tm1688);him-5(e1490)* adult hermaphrodites (n = 20).

To address whether the lack of *daf-7* expression meant that the ASJ cells in *hlh-3(tm1688);him-5(e1490)* adults were not present we examined an ASJ specific *trx-1p::trx-1::gfp* reporter in both genotypes (Sanzo-Machuca et al., 2019). We find that both ASJ cells are detectable in the *him-5(e1490)* and *hlh-3(tm1688);him-5(e1490)* adult males (Figure S2 A-C).

### Mutant male AWA sensory neurons fail to downregulate expression of ODR-10 and show greater attraction to diacetyl

DAF-7/TGF-ß mediated signaling in well fed, adult males has been shown to decrease expression of ODR-10 in adult male AWAs (Wexler et al., 2020). A decrease in ODR-10 has been linked to increased exploration behavior (Ryan et al., 2014). To determine whether ODR-10 expression is altered in *hlh-3(tm1688);him-5(e1490)* adult male AWAs, we measured ODR-10::GFP intensity levels relative to that of a 0.3% InSpeck standard fluorescent intensity bead (Modular Probes). We find that *hlh-3(tm1688);him-5(e1490)* adult male ODR-10::GFP expression was significantly brighter than that in adult *him-5(e1490)* males, but not significantly different than that in *him-5(e1490)* L4 males (Figure 5 A, B, E, and F). *hlh-3(tm1688);him-5(e1490)* adult hermaphrodites ODR-10::GFP levels remain unaffected (Figure 5 C, D, and F)

**Figure 5:**
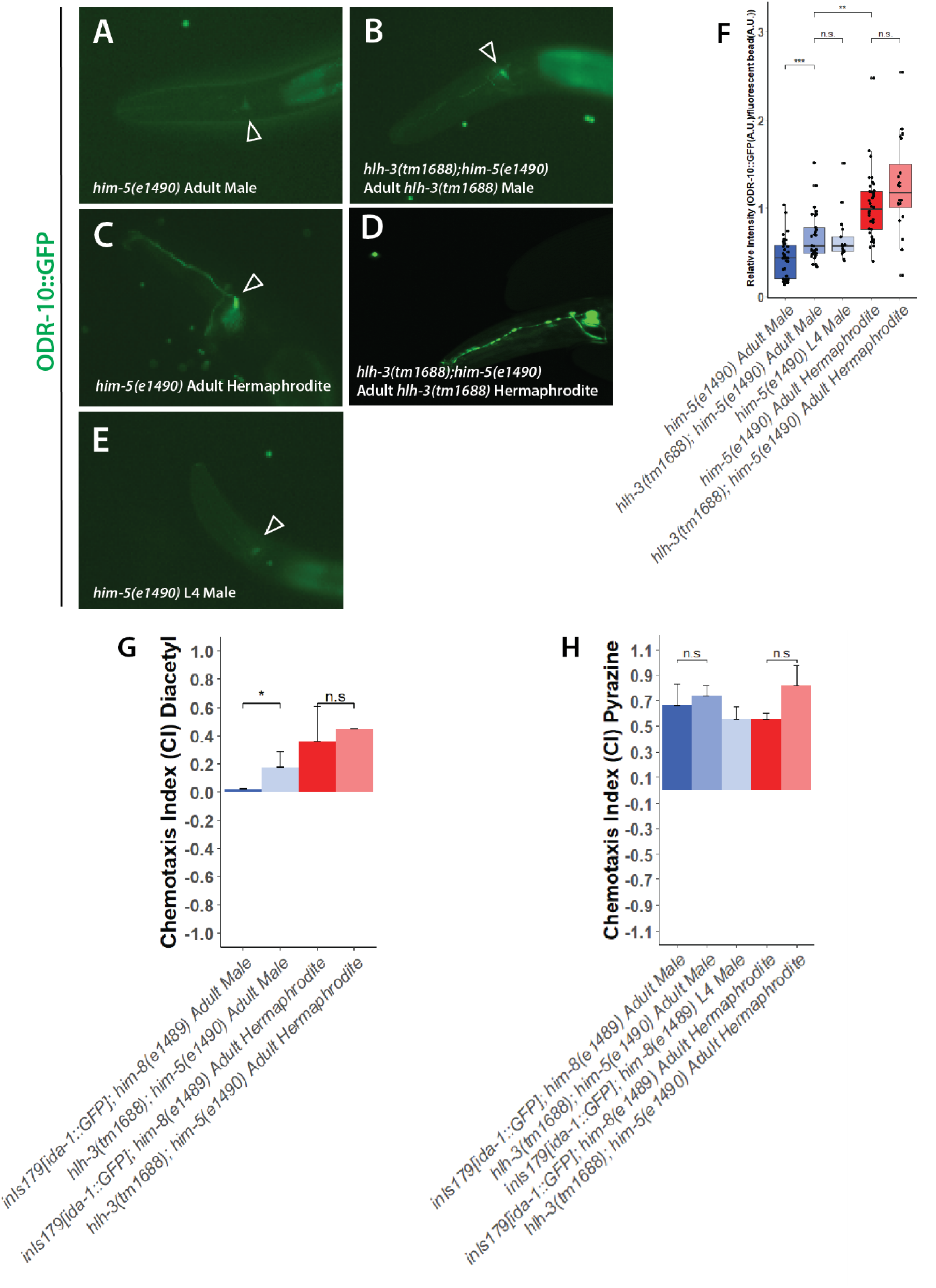
HLH-3 is required for adult male *odr-10* repression and subsequent chemosensation of diacetyl. **(A-E)** Representative images of ODR-10::GFP expression in *him-5(e1490)* adult males, *him-5(e1490)* adult hermaphrodites, *him-5(e1490)* L4 males, and *hlh-3(tm1688);him-5(e1490)* adult males. **(F)** Quantification of ODR-10::GFP expression in relation to 0.3% standard intensity beads for *him-5(e1490)* adult males (n = 39), *hlh-3(tm1688);him-5(e1490)* adult males (n = 30), *hlh-3(tm1688);him-5(e1490)* adult hermaphrodites (n = 20), *him-5(e1490)* L4 males (n = 28), and *him-5(e1490)* adult hermaphrodites (n = 39). *\*\*\* p = 0.000229*, *\*\* p < 0.00001*, *n.s. p = 0.403301, n.s. p value = 0.091906* as determined by Student’s T-test. Expression of ODR-10::GFP is significantly dimmer in *hlh-3(tm1688);him-5(e1490)* adult males than in *him-5(e1490)* adult males. **(G)** Diacetyl (1:1000 dilution) chemotaxis assay where *hlh-3(tm1688);him-5(e1490)* adult males have a mild attraction to diacetyl, while *him-5(e1490)* adult males do not. Values plotted as averages with SEM for 3 independent experiments (n = 40-50 per genotype). **(H)** Pyrazine (0.1mg/1ml) chemotaxis assay where both *him-5(e1490)* and *hlh-3(tm1688);him-5(e1490)* adult males have a strong attraction to pyrazine (n = 40-50 per genotype). Values plotted as averages with SEM for 3 independent experiments (n = 40-50 per genotype).

Others have shown that increased levels of ODR-10 in L4 male AWAs lead to decreased levels of exploration and increased attraction to diacetyl (Colbert & Bargmann, 1995; Ryan et al., 2014). To determine if the observed decrease in ODR-10 expression translates to changes in chemoattraction to diacetyl, we performed a chemotaxis assay (Bargmann et al., 1993; Margie et al., 2013). Consistent with increased ODR-10 expression levels, we found that *hlh-3(tm1688);him-5(e1490)* adult males have a significant increase in attraction to the odorant diacetyl (Figure 5 G). To further determine the functionality of *hlh-3(tm1688);him-5(e1490)* AWA odorant detection, we then assayed attraction to another odorant detected by AWA, pyrazine (Bargmann et al., 1993). Although the receptor to pyrazine is not currently identified, it is known that it is not a ligand of ODR-10 (Sengupta et. al. 1996). Our findings show that *hlh-3(tm1688);him-5(e1490)* adult males and hermaphrodites show chemoattraction of pyrazine similar to that of their *him-5(e1490)* counterparts (Figure 5 H). This indicates that while some chemosensory functions of the AWAs of adult males are abnormal, others remain unaffected.

### Post-embryonic expression of *hlh-3* is detectable in male AIMs, but not AWAs or ASJs

AIMs act non-autonomously to influence the cellular circuitry regulating exploration behavior in adult males (Barrios et al., 2012). AIMs are sex-shared head interneurons and at a position reminiscent of where we find *ic271[hlh-3::gfp]* positive cells (Figure 1 B) in the L1 male. To determine if the identity of the cells present in the head of L1 worms were AIM cells, we introduced *ttx-3p::rfp*, a cell specific reporter for the landmark cell, AIY (Hobert et al., 1997). Given that one of the *ic271[hlh-3::gfp]* positive cells in the head is located immediately anterior to AIY, the landmark cell, we conclude one of the *hlh-3* positive cells in the head are AIMs (Figure 7 A, C). It is noteworthy there is no adjacent *hlh-3* positive cell in L1 hermaphrodites (Figure 7 B, C). *ic271[hlh-3::gfp]* positive AIM cells were not observed in early L3/late L4 stage males or hermaphrodites.

We then determined whether *hlh-3* expression was also detectable in the ASJs and AWAs as well. DiI staining was performed on *ic271[hlh-3::gfp]* L1 and late L3/early L4 staged males to co-label ASJ cells and determine if *hlh-3* was expressed in these cells. There was no evidence that *hlh-3* was present in ASJs at either of these stages (Figure 7 D and E).

To determine if *hlh-3* was expressed in AWA cells, we examined both L1 and late L3/early L4 males co-expressing *podr-10::rfp* and *ic271[hlh-3::gfp]* (Wan et al., 2019). We do not detect *hlh-3 expression* in male AWAs at either stage (Figure 7 F and G).

### Expression of *hlh-3* in male AIMs is not sufficient to down-regulate expression of ODR-10 in AWAs

Since we learned that hat *hlh-3* is expressed post-embryonically in AIM cells, but not AWA or ASJ cells, we hypothesized that a loss of HLH-3 in the AIMs may be non-autonomously causing the phenotypes observed in both other sets of cells. To address whether the function of HLH-3 in the AIM cells is necessary to regulate ODR-10 levels in the AWA cell, we generated a construct in which we utilized a promoter fragment of *eat-4* expressed only in AIMs and PHCs of both males and hermaphrodites to drive expression of a GFP tagged copy of *hlh-3 (Ex[eat-4(prom11):: hlh-3::gfp + rol-6(su1006)])* (Serrano-Saiz et al., 2017). We then measured ODR-10::GFP intensity of this strain in comparison to a control. We find that rescuing expression of HLH-3 in the AIMs of males is not sufficient to rescue ODR-10 levels in the AWAs (Figure 8 A, C, and E).

## DISCUSSION

### Expression of *hlh-3* is sexually dimorphic

The pattern of expression of *hlh-3* in hermaphrodites has been described in the past (Doonan et al., 2008; Krause et al., 1997; Lloret-Fernández et al., 2018; Perez & Alfonso, 2020). However, a description of *hlh-3* expression in males has not been reported before this work. To determine the male expression pattern during post embryonic development, and to ascertain if there were any differences from that in hermaphrodites, we documented the expression patterns of *ic271[hlh-3::gfp]* in both sexes, at all post-embryonic stages, and compared them side by side. As shown before, in hermaphrodites, expression of *hlh-3* was observed in P cells and Pn.a descendants at the L1 stage (Doonan et al, 2008; Perez & Alfonso, 2020). In contrast, the expression pattern in males is expanded to several more cells along the VNC that we suspect to be precursors to the male specific descendants of the P cells, CAs and CPs.

It is also interesting to note that *hlh-3* expression is observed in a few other cells in the head of L1 males, but not hermaphrodites. The location of these cells suggests these positive cells are either the CEMs, or a subset of sex-shared neurons that are necessary for sex specific functions (Shaham & Bargmann, 2002). This work corroborates previous findings that *hlh-3* is expressed in regions of the adult and juvenile staged hermaphrodites. Unfortunately, characterization of the L2 expression pattern in either males or hermaphrodites was not possible because sex discrimination at this stage, using our equipment, is not feasible.

Expression of *hlh-3* at the L3 stage and beyond differs in males when compared to hermaphrodites as it is observed in a much higher number of cells in the tail. Given that HLH-3 is required for the terminal differentiation of both VCs and HSNs, it is reasonable to assume that a subset of the *hlh-3* positive cells observed in male tails are also sex-specific (Doonan et al., 2008; Krause et al., 1997; Perez & Alfonso, 2020). Because of its location, the cluster of expression observed in the tail ganglia is likely to include neurons that play major roles in male mating and mate sensing activities, such as ray A-type or B-type neurons (RnA and RnB), hook sensillum neurons (HOA and HOB), or phasmid neurons (PHA, PHB, and PHC) (Barr & Sternberg, 1999; Garcia et al., 2001; K. S. Liu & Sternberg, 1995; Loer & Kenyon, 1993). Efforts to utilize the whole nervous system NeuroPal reporter to positively identify each of the cells that express *hlh-*3 were not fruitful because nuclear expression of the CRISPR-Cas9 *ic271[hlh-3::gfp]* allele was dim (Tekieli et al., 2021). Further investigation with an alternative *hlh-3* reporter is required to confirm the identity of the *hlh-3* positive cells in the male.

Post-embryonic expression of *hlh-3* in males is most prominent in the L1 stage as well as the late L3/early L4 stages in a sexually dimorphic pattern. Previous reports show sexually dimorphic neurons display ‘just-in-time’ development, in which neurons that acquire sexually dimorphic terminal identity features form these features in the L4 stage just before adulthood (Hobert, 2016; Tekieli et al., 2021). The expression pattern of *hlh-3* at the L3/L4 stages and in places such as the male tail support the idea that *hlh-3* function is necessary for many male specific neurons to mature to adulthood; this is consistent with the hypothesis that it is necessary for the terminal differentiation of hermaphrodite specific neurons (Doonan et. al 2008; Perez & Alfonso 2020). However, the expression of *hlh-3* in some cells in the male L1 stage, but not later stages, suggests that *hlh-3* may be necessary to set into motion a cascade of events that contribute to terminal identity acquisition at a later stage. These contrasting patterns of expression suggest that the function of *hlh-3* is necessary at different developmental stages to promote neuronal identity cascades in motion.

### Many neurons necessary for male exploration fail to obtain terminal differentiation features

Our results have shown that the sex-shared AWAs, ASJs, and AIMs of adult mutant males fail to acquire terminal identity features, and subsequently do not result in a defect in male exploration. Mutant adult male AIM neurons fail to fully acquire the mature cholinergic identity (Figure 3). In turn, male ASJ neurons are present but fail to upregulate expression of *daf-7* upon adulthood (Figure 4). As a result, expression of ODR-10 is not downregulated in adult male AWA neurons, resulting in increased attraction to diacetyl (Figure 5).

Since *hlh-3* is not expressed post-embryonically in either the AWAs or ASJs (Figure 6), we asked whether restoration of *hlh-3* function in AIM, the only cell in the circuit where we see post-embryonic expression of *hlh-3*, could restore function to the circuit in a non-cell autonomous way. We find it does not (Figure 7). Expression of *hlh-3* in the L1 to L4 stages does not rescue the mutant phenotypes in the AWA cell and by inference the ASJ cell. This is surprising given that the current body of evidence suggest that the emergence of male sexual dimorphism results from “just-in-time” development (Tekieli et al., 2021); a process in which dimorphic neurons are born in any given stage, but all begin the terminal differentiation program in the L4 stage (Tekieli et al., 2021). Rescue of ODR-10 expression levels in the AWAs through rescue of *hlh-3* expression in AIMs may be due to one of a few reasons.

**Figure 6:**
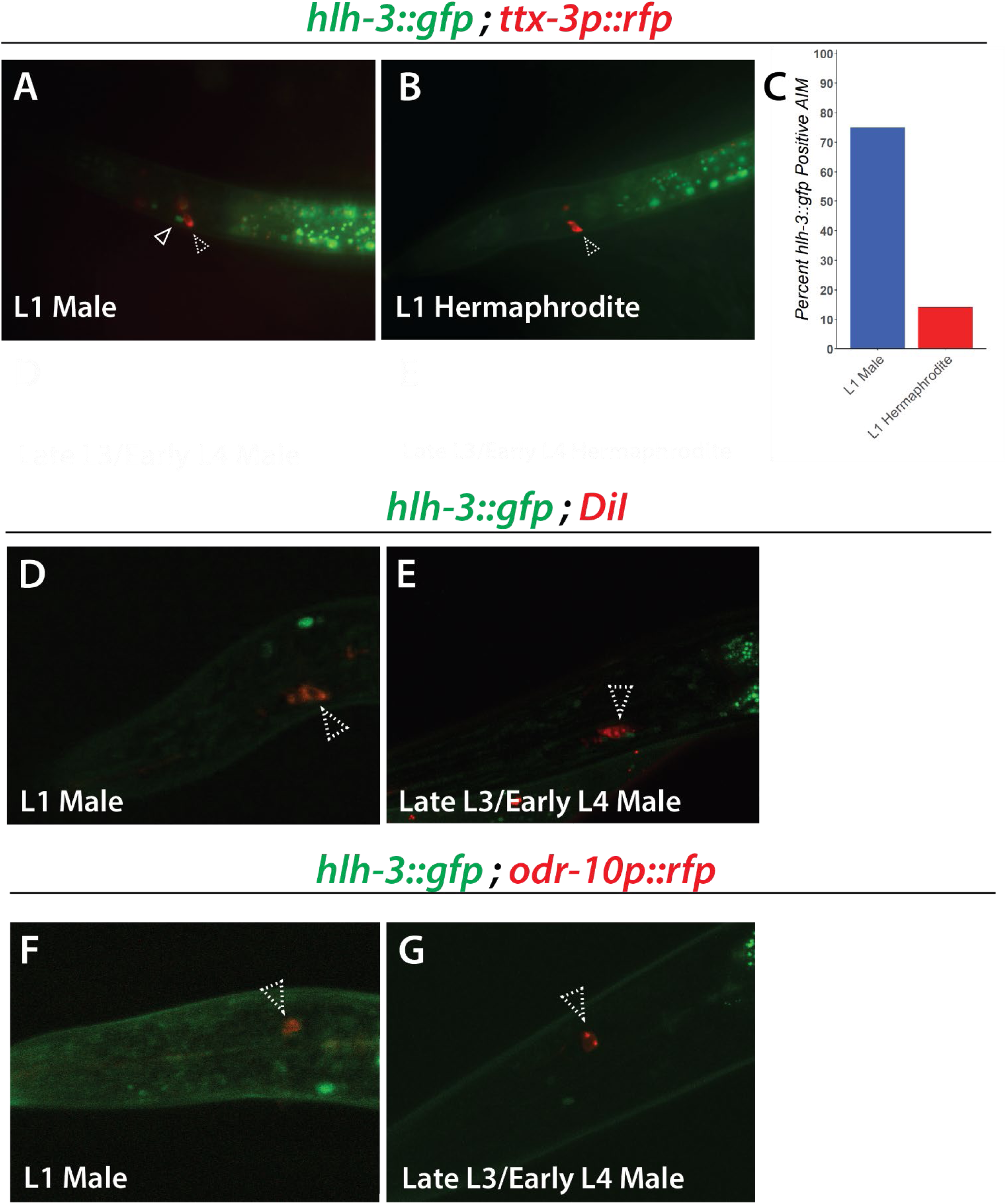
*hlh-3* is expressed post-embryonically in male L1 AIMs. **(A-B)** Representative images of *ic271[hlh-3::gfp]* and *ttx-3p::rfp* expression in the head of an L1 male and hermaphrodite, respectively. **(C)** Quantification of *ic271[hlh-3::gfp]* expression in the AIM cell of L1 males (n = 25) and hermaphrodites (n = 23). Open arrowheads indicate AIM cells, dashed arrowheads indicate AIY cells. **(D-E)** Representative images of *ic271[hlh-3::gfp]* and *ttx-3p::rfp* expression in the head of a late L3/earlyL4 male and hermaphrodite, respectively. (F-G) Representative images of *ic271[hlh-3::gfp]* and DiI stain in the head of an L1 and late L3/earlyL4 male, respectively. Dashed arrowhead represents the ASJ cells. (H-I) Representative images of *ic271[hlh-3::gfp]* and *odr-10p::rfp* expression in the head of an L1 and late L3/earlyL4 male, respectively. Dashed arrowhead indicates AWA cells.

First, the *eat-4(prom11)::gfp* construct used to drive expression of *hlh-3* in AIMs was not integrated. While expression of the reporter is present and detectable in hermaphrodites of all stages, in males, expression is only observed in the L1 through L4 stages. Assessment of L4 stage males shows the construct present in 46% of L4 worms, when the construct is still detectable. Thus, it is possible that in some assayed males the construct was lost in precursors to the AIM. Since the rescue construct is not expressed in adult males, choosing adult males in which it is certain that the construct is present was impossible. Thus, the lack of rescue could be explained by a skewed proportion of males in which the extra chromosomal array is lost in the AIM cells.

Secondly, it could be suggested that the construct is unable to generate functional *hlh-3* products. However, lack of function of the HLH-3 protein produced by the rescue construct is unlikely. The *eat-4* construct was generated from a *punc-86::hlh-3::gfp* construct that has been shown to rescue HSN maturation defects in *hlh-3(tm1688)* adult hermaphrodites (Raut, 2017). Moreover, the fact that we detect expression of the *eat-4(prom11)::gfp* construct in hermaphrodites suggests it has been transcribed and translated.

One other possibility is that some other neuron influencing this circuitry is required for the down-regulation of ODR-10 in the AWAs. The network of neurons characterized in this study does not extend to all neurons that regulate male exploration behavior (Barrios, 2012). It is possible that other neurons in the head (URY) or tail (PHA and PQR), that have been shown to affect male exploration, may require expression of *hlh-3* in order to repress ODR-10 in adult males. This would require further work to characterize their differentiation and functionality within *hlh-3(lof)* males.

Lastly, it is possible that to rescue the mutant phenotype, it is necessary to restore *hlh-3* function in embryonic stages. It is known that *hlh-3* is expressed transiently in embryonic AIM, AWA, and ASJ cells of hermaphrodites (Ma et. al., 2021). Thus, it is possible that *hlh-3* function is necessary embryonically in males to set in motion a regulatory cascade for one or all of these cell types. If so, our observations are, in the context of the AIM, inconsistent with the hypothesis of “just-in-time” maturation of sex-shared neurons with sexually dimorphic functions. To further explore the mechanism of *hlh-3* function it would be necessary to determine whether *hlh-3* expression is also detectable in male AIM, AWA, and ASJ neurons in the embryonic stage and a rescue construct driving embryonic expression would need to be generated.

From this data we can conclude that *hlh-3* expression is necessary for this subset of neurons to acquire terminal identity features, though further work is necessary to determine the mechanism by which that is accomplished.

### *hlh-3* is a non-traditional bHLH protein with a late acting role in neuronal terminal identity acquisition of sex-specific and sex-shared neurons with sex-specific functions

In this work we have characterized the phenotypes displayed by *hlh-3(lof)* adult male neurons that are sex-shared but have sex-specific roles in adulthood. Our findings regarding defects in sex shared neurons with sex-specific functions in *hlh-3* mutant males adds support to the hypothesis that *hlh-3* is necessary for the terminal differentiation of both sex-specific and sex-shared neurons involved in sex-specific behaviors, a novel finding. How HLH-3 regulates these processes is not yet clear. However, we can say that the role of *hlh-3* is not that of a traditional proneural bHLH transcription factor but rather one of the many bHLH proteins in *C. elegans* that has non-traditional roles in development. Unlike what has been observed in the hermaphrodite specific HSNs and VCs, in males *hlh-3* expression may not necessary throughout the remainder of developmental stages. This indicates that *hlh-3* may be acting much earlier in development to set up a signaling cascade that leads to the differentiation of these neurons at a much later stage than previously hypothesized. Alternatively, it may be possible that our reporters are not able to detect minute quantities of expressed *hlh-3* that may be required for the acquisition of these terminal differentiation features. In this scenario, if transcripts were detectable in the late L3/early L4 stage then this would much better fit the current model that indicates that many factors that contribute to the arising of sexually dimorphic features are expressed just before these changes are observed to occur.

If embryonic expression of *hlh-3* does truly set up a signaling cascade that results in attainment of sexually dimorphic features emerging in the L4 to adult transition, then it is possible that the process of sex-shared neurons gaining these sexually dimorphic features may start much earlier than previously thought. Early acting factors, such as *hlh-3* and/or other bHLH proteins, may be necessary to specify sex-shared neurons with the potential to acquire these sexually dimorphic features earlier in development, but then factors such as *mab-3* and *lin-29a* may be functioning in parallel, late acting pathways to more directly carry out acquisition of these features. Re-emergence of *hlh-3* expression in a subset of these neurons may be indicative of re-enforcement of the identity of these cells. Further work deciphering the early acting, embryonic roles of *hlh-3* versus the late acting, post-embryonic role of *hlh-3* in sex-shared neurons through use of condition specific knockout strains would answer questions concerning this topic.

## MATERIALS AND METHODS

### Strain maintenance

All strains were maintained at 22°C under standard conditions on nematode growth media seeded with OP50-1 (Brenner, 1974; Johnson et al., 1988). The National Bioresource of Japan generated the *hlh-3(tm1688)* allele used in this work. Several strains were provided by the CGC, which is funded by NIH Office of Research Infrastructure Programs (P40 OD010440). VZ797 was graciously provided by Dr. Antonio Miranda-Vizuete. *podr-10::rfp* constructs were graciously provided by Dr. King L Chow.

### Construction of transgenic strains

The genomic region of chromosome III contained between -2609bp and -2155bp from the ATG of the *eat-4* locus was amplified via PCR (Serrano-Saiz et al., 2017). The DNA fragment was cloned into the pRD6 construct after excising the *unc-86* promoter with *SalI* and *HindIII* enzymes to generate pKA1 (Raut, 2017). pKA1 (50ng/µl) was injected into *hlh-3(tm1688) II; kyIs37[ODR-10::GFP + lin15(+)] II; him-5(e1490) V* (AL190) using *rol-6(su1006)* (50ng/µl) as a co-injection marker to generate *[eat-4(prom11)::hlh-3::gfp + rol-6(su1006)]; hlh-3(tm1688) II; kyIs37[ODR-10::GFP + lin15(+)] II; him-5(e1490) V* (Peixoto et al., 1998). This produced strains in which *hlh-3* function was only rescued in AIMs and PHCs of both males and hermaphrodites.

*podr-10::rfp* (50ng/ul) was injected into *ic271[hlh-3::gfp]; him-5(e1490)* using *rol-6(su1006)* (50ng/µl) as a co-injection marker (Peixoto et al., 1998; Perez & Alfonso, 2020; Wan et al., 2019). This generated *[podr-10::rfp + rol-6(su1006)]; ic271[hlh-3::gfp]; him-5(e1490) V*. This produced a strain which co-labels *ic271[hlh-3::gfp]* and *podr10::rfp* expressing cells.

### Microscopy

#### Fluorescent Imaging

Animals were segregated by sex at the L4 stage on uncrowded plated seeded with a lawn of OP50-1. After 26 hours they were mounted on 2.5% agarose pads and anesthetized with 10mM levamisole (Johnson et al., 1988). Images were taken at either 63x or 100x with a Carl Zeiss Axioskop 2 fluorescence microscope utilizing Axiovision 2.002 software. Images were further visualized with Fiji version 1.54f software (Schindelin et al 2012). Original grayscale images were converted with red and green Lookup tables and merged using Merge Channels. Figures were created using Adobe Illustrator version 28.6.

#### Fluorescence Quantification

GFP fluorescence was quantified by measuring the intensity of the reporter using Fiji software and comparing it to an InSpeck™ Green (505/515) microscope image of a 0.3% standard intensity bead mounted on the same agarose pad (Invitrogen, I7219, Schindelin et al., 2012). Statistical significance was determined using the Student’s T-test. About 30-40 cells were imaged per genotype.

Quantification of *cho-1p::rfp* intensity in AIM cells was performed by comparing its intensity relative to that of the *cho-1p::rfp* intensity in the neighboring AIY cells (Serrano-Saiz, Pereira, et al., 2017). Statistical significance was determined using the Student’s T-test. About 25-35 cells were imaged per genotype.

### Leaving assay

L4 males or hermaphrodites were picked 26 hours prior to the assay and isolated on an uncrowded plate seeded with OP50-1 bacteria. Exploration assays were conducted as previously described (Lipton et al., 2004). A single animal was placed on a 10cm plate seeded with 20µl of OP50-1 bacteria. Animals were allowed to roam for 1 hour before their tracks were observed. If the animal reaches an area 1cm from the edge of the plate (the scoring distance), they are considered a “leaver”. The number of leavers is counted at the end of each hour time point for a total of 5 hours. The probability of leaving was determined by obtaining the hazard value (λ), using the single exponential decay function N(t)/N(0)=exp(-λt). These values were calculated using the exponential parametric survival using maximum likelihood (R Project for Statistical Computing with R survival package). Statistical significance was calculated using one-way ANOVA with Tukey HSD.

For the locomotion control, 10cm plates were seeded with OP50-1 bacteria spread into a lawn covering the area to the 1cm scoring distance used in the exploration assay (Barrios et al., 2012). Single animals were placed on these plates and their tracks were analyzed using the same methodology as the exploration assay. Statistical significance was calculated using chi-squared analysis.

### Chemotaxis assays

About 50-60 L4 males were picked 26 hours prior to the assay and isolated on an uncrowded plate seeded with OP50-1 bacteria. To minimize transfer of bacteria, adult males were transferred to an unseeded plate before being washed off with M9 (Stiernagle, 2006).and collected in a 1.5mL tube. Worms were then spun down and transferred to the assay plate. The rest of the assay was conducted as previously described (Margie et al., 2013). 2µl of diacetyl (1:1000µl dilution in ethanol) was used in diacetyl assays (Arcos Organics, 431-03-8). 2µl of a 1:100µl dilution of 10mg/ml pyrazine in ethanol was used in pyrazine assays (Oakwood Chemical, 290-37-9). For each chemotaxis assay, averages are shown pooled across 3 replicates (n = 40-50 per genotype). Significance was determined using the Student’s T-test.

### Data Availability

Strains and plasmids are available upon request. The authors affirm that all data necessary for confirming the conclusions of the article are present within the article, figures, and tables.

## ACKNOWLEDGEMENTS

We would like to acknowledge Maryam Sabir for help with data acquisition; Dr. Suzanne McCutcheon and Dr. Janet Richmond for imaging equipment; and the Department of Biological Sciences for support with this publication. We thank Dr. Antonio Miranda-Vizuete and Dr. King L Chow for graciously providing strains and constructs.

---

**Figure S1:**
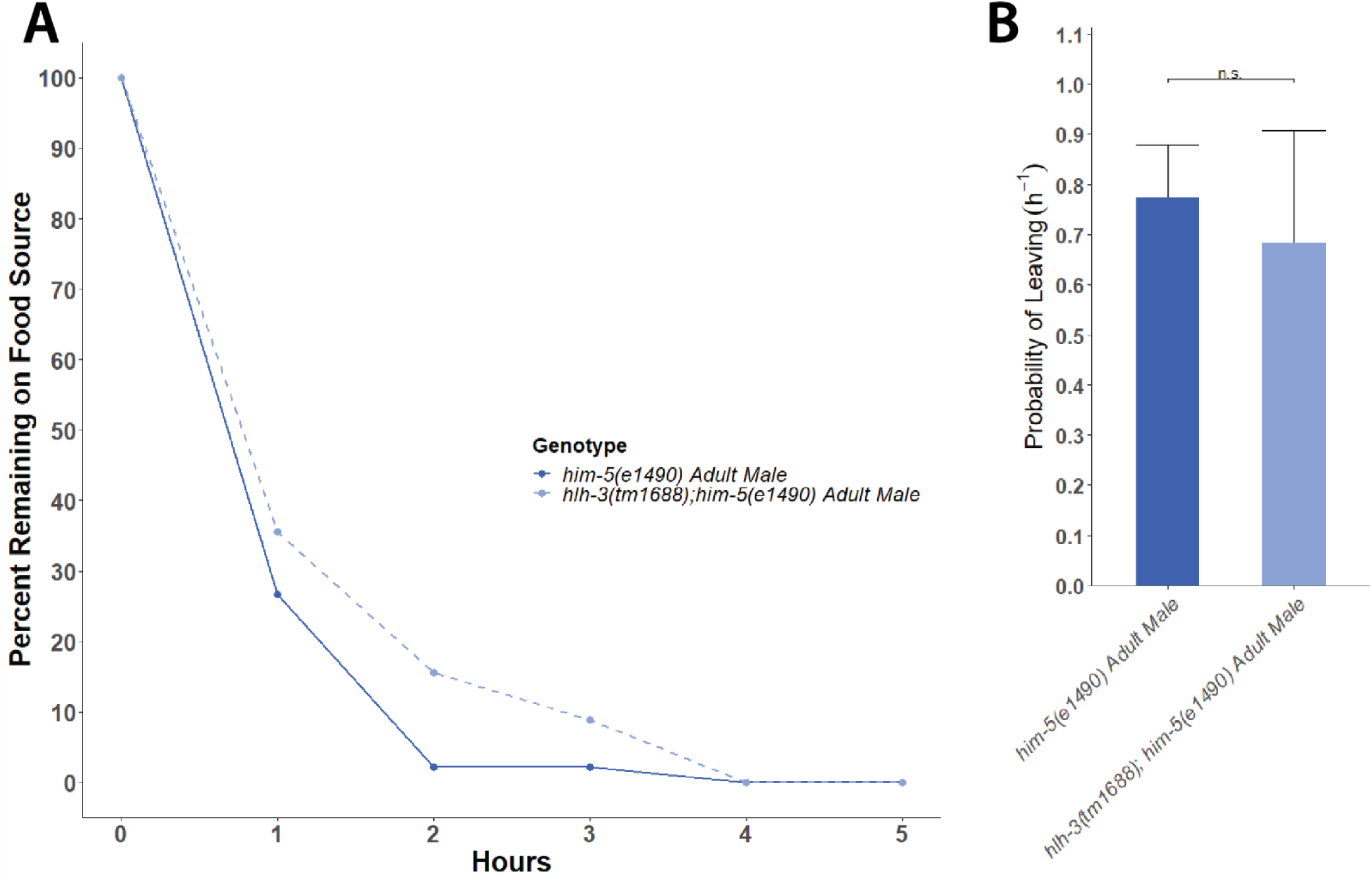
HLH-3 null adult males can reach scoring distances at similar rates as wild type males. **(A)** Representative graph depicting the quantification of average male specific exploration behavior of *him-5(e1490)* adult males and *hlh-3(tm1688);him-5(e1490)* adult males (n = 45 for each genotype). Each point represents the percentage of worms remaining in the non-leaver area at the indicated time point. **(B)** Probability of leaving a food source for *him-5(e1490)* males and *hlh-3(tm1688);him-5(e1490)* adult males was calculated by the single exponential decay function N(t)/N(0)=exp(-λt), where N(t)/N(0) refers to the ratio of non-leavers to leavers at a given time (t), and the hazard value (λ) is the constant rate demonstrating the probability of leaving. Values plotted as averages with SEM for 3 independent experiments. *n.s. p = 0.961256* as determined by chi-squared analysis.

**Figure S2:**
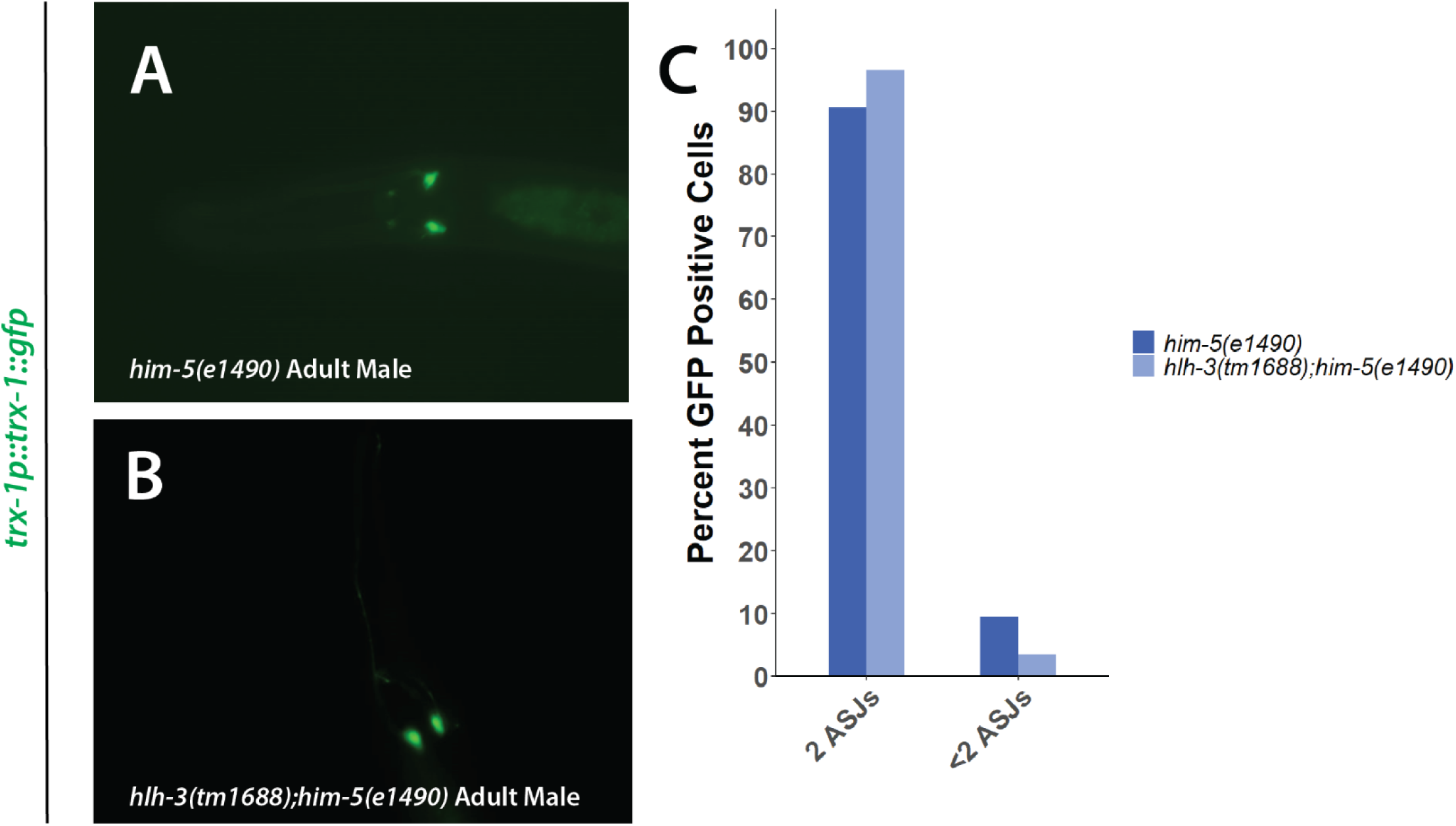
Adult male ASJs are detectable in wild type and mutant adult males; they still express *trx-1*. **(A-B)** Representative images of *trx-1p::trx-1::gfp* expression in adult male *him-5(e1490)* and *him-5(e1490);hlh-3(tm1688)*, respectively. **(C)** Quantification of indicated genotype containing either 2 or <2 ASJ cells for *him-5(e1490)* adult males (n = 21) and *him-5(e1490);hlh-3(tm1688)* adult males (n = 29).

