## Supplement for "HLH-3 Is Required for the Maturation of Neurons Necessary for Male Specific Exploration Behavior"

**RUNNING TITLE:**

*hlh-3* is required for the differentiation of neurons required for male specific exploration behavior

**KEYWORDS**

*hlh-3*, differentiation, AIM, ASJ, AWA, sexually dimorphic

**CORRESPONDING AUTHOR INFORMATION**

Aixa Alfonso

University of Illinois at Chicago

840 W. Taylor, M/C 067

Chicago, IL 60607

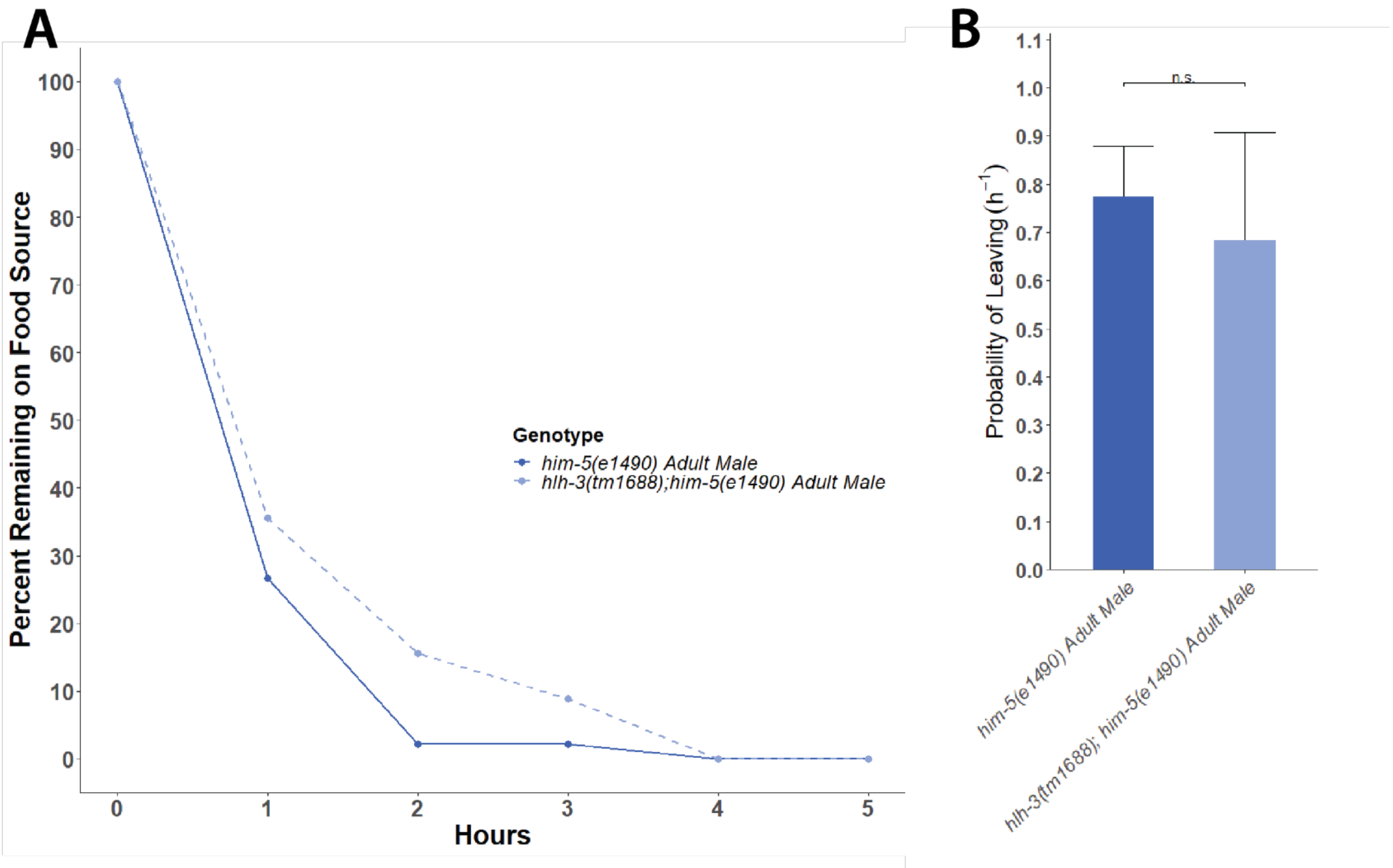


Figure S1: HLH-3 null adult males can reach scoring distances at similar rates as wild type males

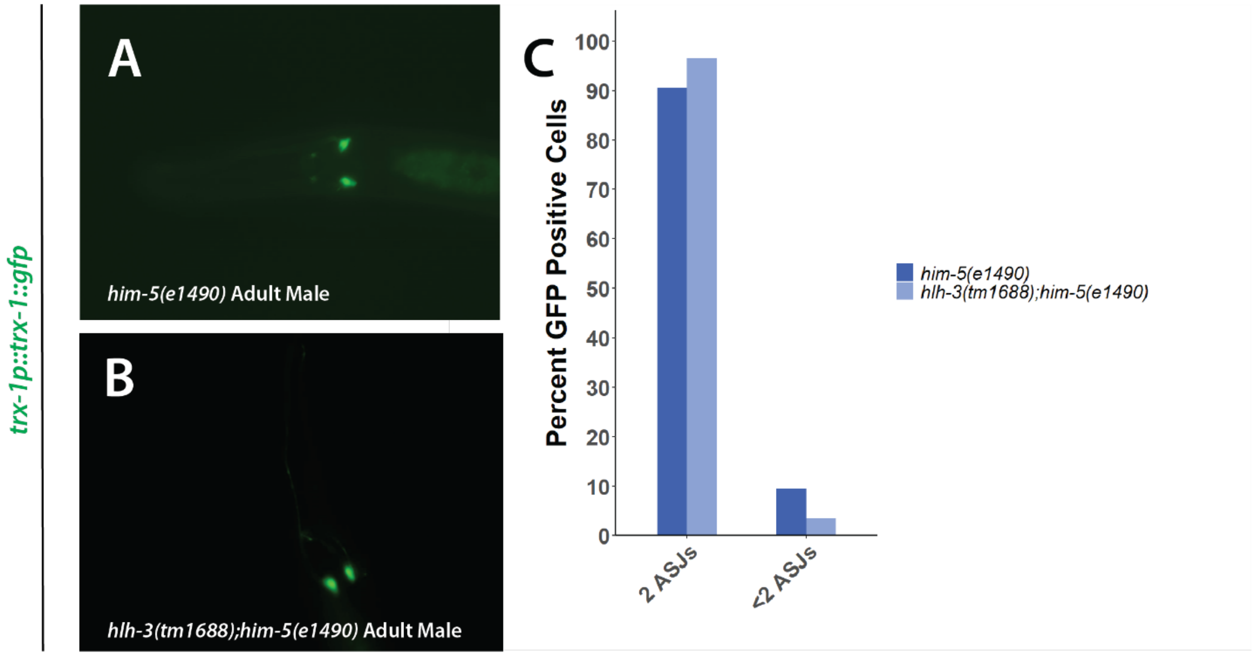


Figure S2: Adult male ASJs are detectable in wild type and mutant adult males; they still express *trx-1*

| **Strain Name** | **Genotype** |
|---|---|
| AL186 | *kyIs37[ODR-10::GFP + lin-15(+)] II; him-5(e1490) V* |
| AL190 | *hlh-3(tm1688) II; kyIs37[ODR-10::GFP + lin15(+)] II; him-5(e1490) V* |
| AL274 | *hlh-3(tm1688) II; ksIs [daf-7p::GFP + rol-6(su1006)]; him-5(e1490) V* |
| AL277 | *ksIs2[daf-7p::GFP + rol-6(su1006)]; him-5(e1490) V* |
| AL299 | *hlh-3(tm1688) II; otIs1[pBHL98(lin-15ab+); ptrx-1::trx-1::GFP]; him-5(e1490) V* |
| AL336 | *hlh-3(tm1688) II; him-5(e1490) V* |
| AL353 | *hlh-3(tm1688) II; otIs544 [cho-1(fosmid)::SL2::mCherry::H2B + pha-1(+)]; mgIs18 [ttx-3p::GFP] IVI; him-5(e1490) V* |
| AL354 | *otIs544 [cho-1(fosmid)::SL2::mCherry::H2B + pha-1(+)]; mgIs18 [ttx-3p::GFP] IV; him-5(e1490) V* |
| AL355 | *hlh-3(ic271[ic271[hlh-3::gfp]]); him-5(e1490) V; otIs133[pttx-3::RFP + unc-4(+)]* |
| AL367 | *hlh-3(ic271[ic271[hlh-3::gfp]]); him-5(e1490) V* |
| AL375 | *ExIs376[eat-4(prom11)::hlh-3::gfp +rol-6(su1006)]; hlh-3(tm1688) II; kyIs37[ODR-10::GFP + lin-15(+)] II; him-5(e1490)V strain 1* |
| AL376 | *ExIs376[rol-6(su1006)]; hlh-3(tm1688) II; kyIs37[ODR-10::GFP + lin-15(+)] II; him-5(e1490)V* |
| AL379 | *ExIs379[podr-10::rfp + rol-6(su1006)]; ic271[hlh-3::gfp]; him-5(e1490) V* |
| BL5715 | *inIs179[ida-1p::GFP] II; him-8(e1489) IV* |
| DR466 | *him-5(e1490) V* |
| KP4 | *glr-1(n2461)III* |
| OH14018 | *otIs520 [eat-4(prom11)::GFP + ttx-3::mCherry]; him-5(e1490) V* |
| PT8 | *pkd-2(sy606) IV; him-5(e1490) V* |
| PT621 | *myIs4 [PKD-2::GFP+Punc-122::GFP] V; him-5(e1490) V* |
| PT2248 | *pdf-1(tm1996) III; him-5(1490) V* |
| VZ797 | *otIs1 [pBHL98(lin-15ab+); ptrx-1::trx-1::GFP]; him-5(e1490) V* |

Supplementary Table 1: Strain list
